# Discovery of an NRAS isoform and activation-state selective macrocyclic peptide

**DOI:** 10.64898/2026.07.30.741788

**Authors:** Kenneth K. Hallenbeck, Yunpeng Zhou, Hubert Josien, Songnian Lin, Aileen Soriano, Todd Mayhood, Jaclyn Robustelli, Xiaomei Chai, My Sam Mansueto, Gireedhar Venkatachalam, Ryan E. Loy, Peng-Hsun Chase Chen, Huifang Yao, Haihong Zhou, Elsa Beyer Krall, David G. McLaren, Adam Weinglass, S. Adrian Saldanha

## Abstract

Macrocyclic peptides have gained increased attention amid claims they are a “Goldilocks” therapeutic modality that can encode the selectivity of a biologic in a footprint close to that of a small molecule. Here we attempt to find a peptide that binds selectively to NRAS, sparing HRAS and KRAS, while accessing the cytosol via passive cell permeability. To do so, we combine subtractive affinity selection with mRNA display to identify Compound 1, an 11mer macrocyclic peptide which binds NRAS at a novel allosteric site between Helix 3 and Helix 4 of the GTPase domain. Compound 1 has total isoform selectivity and can be tuned to achieve activation-state selectivity with a single amino acid change. While it has preferential affinity for oncogenic NRAS-specific mutations, it does not inhibit NRAS function or achieve passive membrane permeability.

## Introduction

RAS activation is implicated in ~20% of all human cancers [1] and intense investment has produced the first therapies targeting the family [2, 3]. However, these drugs target oncogenic mutations in KRAS and cannot treat the full spectrum of RAS-driven cancers. Hotspot activating mutations in NRAS occur in 3% of all tumors across several tumor types [1]. Genetic dependency on NRAS is highly associated with NRAS hotspot mutations *in vitro*, with most NRAS wildtype cell lines showing minimal viability effect when NRAS is knocked out [4]. In addition, NRAS knockout mice are viable and fertile with normal development, while KRAS knockout is embryonic lethal [5]. These preclinical observations suggest that selectively targeting NRAS in NRAS-mutant tumors may provide a better therapeutic index than pan-RAS inhibitors under development [6,7].

Unfortunately, the high sequence similarity of KRAS, HRAS and NRAS (>90% in the structured regions) has confounded the development of isoform-specific inhibitors. To date, RAS isoform selectivity has been achieved by targeting oncogenic mutants with small molecules [2,3,8–10], other sites with proteins such as monobodies [11] or de novo designs against the hyper variable region at the C-terminus [12].

One cell-permeable macrocyclic peptide has been reported to bind RAS proteins: LUNA-18. It is a pan-RAS inhibitor identified with mRNA display [13], a technology which encodes libraries of over a trillion unique peptides. Such diversity may be large enough to achieve high affinity and selectivity in scaffolds small enough to have passive cell permeability [14–16].

To identify a RAS isoform-specific macrocyclic peptide, we combine mRNA display with an affinity selection strategy that first subtracts all KRAS binders from the displayed library, then attempts to enrich for NRAS binders from the remaining members. The result is Compound 1, a highly selective 11mer peptide that binds to a cleft between NRAS Helix 3 and Helix 4, taking advantage of four NRAS-specific amino acids to achieve total selectivity over KRAS and HRAS.

## Results and Discussion

A 9-12mer mRNA display library, randomized and containing 5 non-natural amino acids and 10 natural amino acids (Fig S1), was screened against an NRAS construct (2-172) containing only the structured domain, since the C-terminal hypervariable domain is post-translationally modified by farnesylation and palmitoylation to enable membrane insertion [17]. The affinity selection proceeded against streptavidin beads loaded with biotinylated NRAS for 1 round, then beginning at round 2, a subtraction step using streptavidin beads loaded with biotinylated KRAS (2-169) was inserted prior to exposure to NRAS (Fig 1A). After a total of 5 rounds of affinity selection, the resulting library was analyzed with Next Generation Sequencing to identify enriched peptides. One enriched peptide was Compound 1, a thioether macrocycle with the sequence ClAc-F-D-Tic-Y-NMeF-Sar-Bip-P-Y-N-C-G (Fig 1B). Compound 1 was synthesized with Solid Phase Peptide Synthesis and tested with Surface Plasmon Resonance (SPR) for affinity to the inactive state of RAS in the presence of GDP. Compound 1 has a *K*_D_ of 1670 nM for NRAS-GDP (Fig 1C) and no detectable affinity for KRAS-GDP and HRAS-GDP (Fig 1D, 1F). To confirm Compound 1 retained affinity for oncogenic NRAS mutations, we also measured affinity for NRAS Q61R-GTP and Q61K-GTP (Fig 1E). Interestingly, Compound 1 has >15-fold higher affinity for the activated mutants loaded with GTP than WT NRAS loaded with GDP, suggesting the pocket it engages is allosterically coupled with NRAS activity.

**Figure 1:**
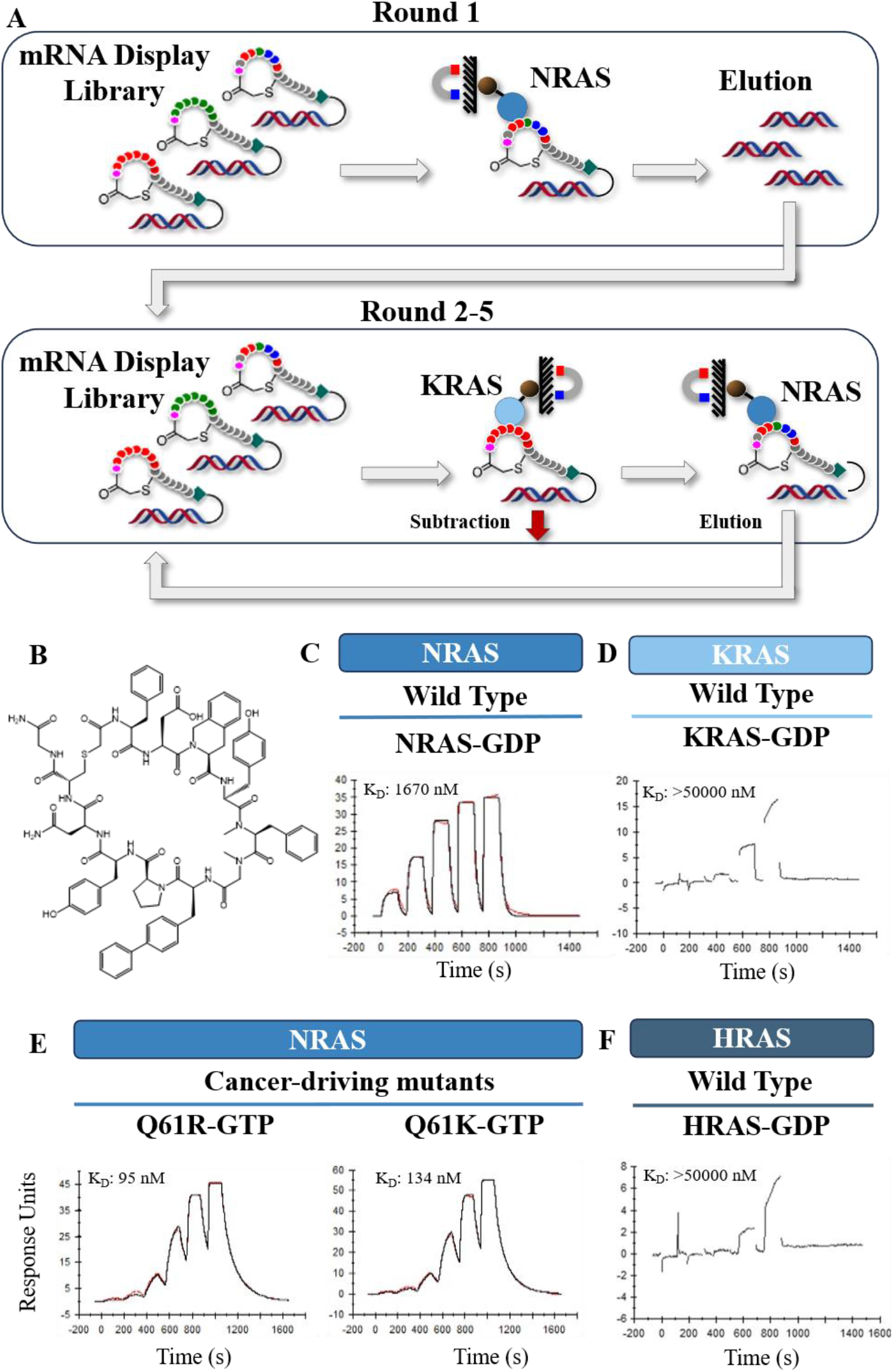
mRNA Display identifies Compound 1. A) Affinity selection strategy. B) Structure of Compound 1. C) SPR Sensorgram for Compound 1 binding to WT NRAS-GDP. D) SPR Sensorgram for Compound 1 binding to WT KRAS-GDP. E) SPR Sensorgram for Compound 1 binding to two NRAS-GTP cancer mutants. E) SPR Sensorgram for Compound 1 binding to WT HRAS-GDP.

To understand the mechanism of this coupling, as well as Compound 1’s isoform selectivity, we mapped the peptide binding epitope by comparing the ^1^H/^15^N HSQC Spectra of NRAS-GDP in presence and absence of Compound 1 (Fig 2A). Comparing the observed chemical shift perturbations (CSPs) to the 3D structure of NRAS previously determined by X-Ray Crystallography [18] (Fig 2B) showed Compound 1 binding induced the most CSPs in Helix 3 and Helix 4 of the GTPase domain (Fig 2C), suggesting the peptide binds NRAS at the cleft between the two helices. This cleft contains four residues which are specific to NRAS, S87, A91, N94 and E132, explaining Compound 1’s inability to bind KRAS and HRAS. In addition, moderate CSPs are observed on Helix 2 of Switch II, which is in direct contact with the cofactor of GTP or GDP. This suggested peptide binding and the NRas active states are coupled through the conformational change of Switch II. Given the distance of the Compound 1 epitope from the GTPase active site and the CRAF protein-protein interaction site, we anticipated it would not modulate NRAS activity. We confirmed this lack of activity in a GNE-SOS ATP transfer assay (Fig 2D) as well as in a NRAS:CRAF TR-FRET PPI assay (Fig 2E). We endeavored to optimize the NRAS affinity of Compound 1 with a series of single amino acid mutations (Table 1), hoping that increased affinity might lead to measurable allosteric inhibition. At position 2, closing the Asp side-chain to form a pyrrolidone ring increased affinity 2.5-fold (Compound 6). Altering position 6 from Sarcosine to d-Proline led to a 3-fold increase in affinity, presumably from the stabilization of the binding conformation. However, when these two changes were combined, the benefits were lost (Compound 23). Changes at position 1 and 9 also provided modest affinity increases. Particularly exciting was the ability to remove a charge from the Asp at position 2 (Compound 6, D to PyD), and to shrink the side chains at position 1 (Compound 2, F to Nle) and position 9 (Compound 19, Y to F) (Table 1, Table S2).

**Table 1:**
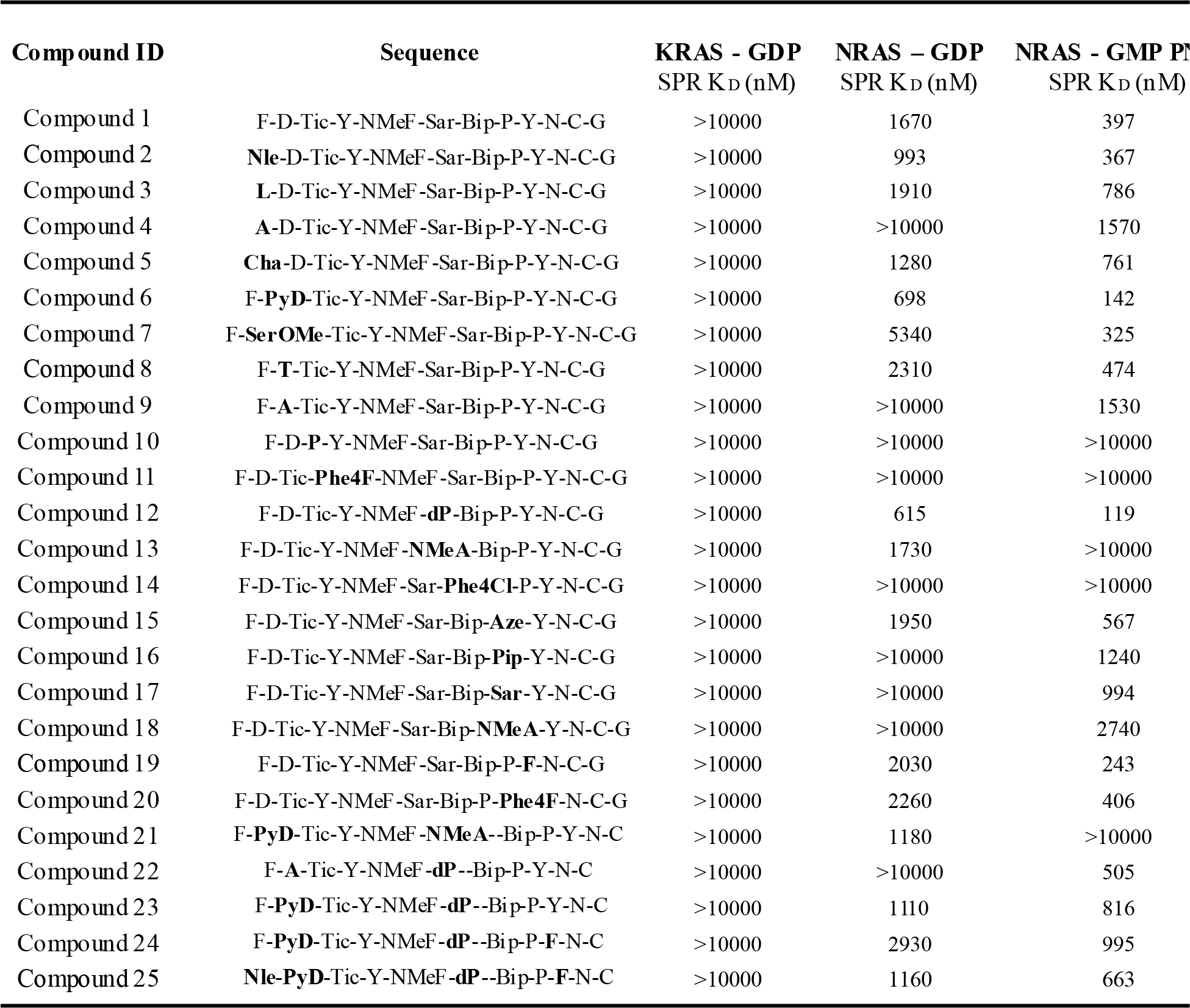
S R Affinity of Compound 1 and Variants.

**Figure 2:**
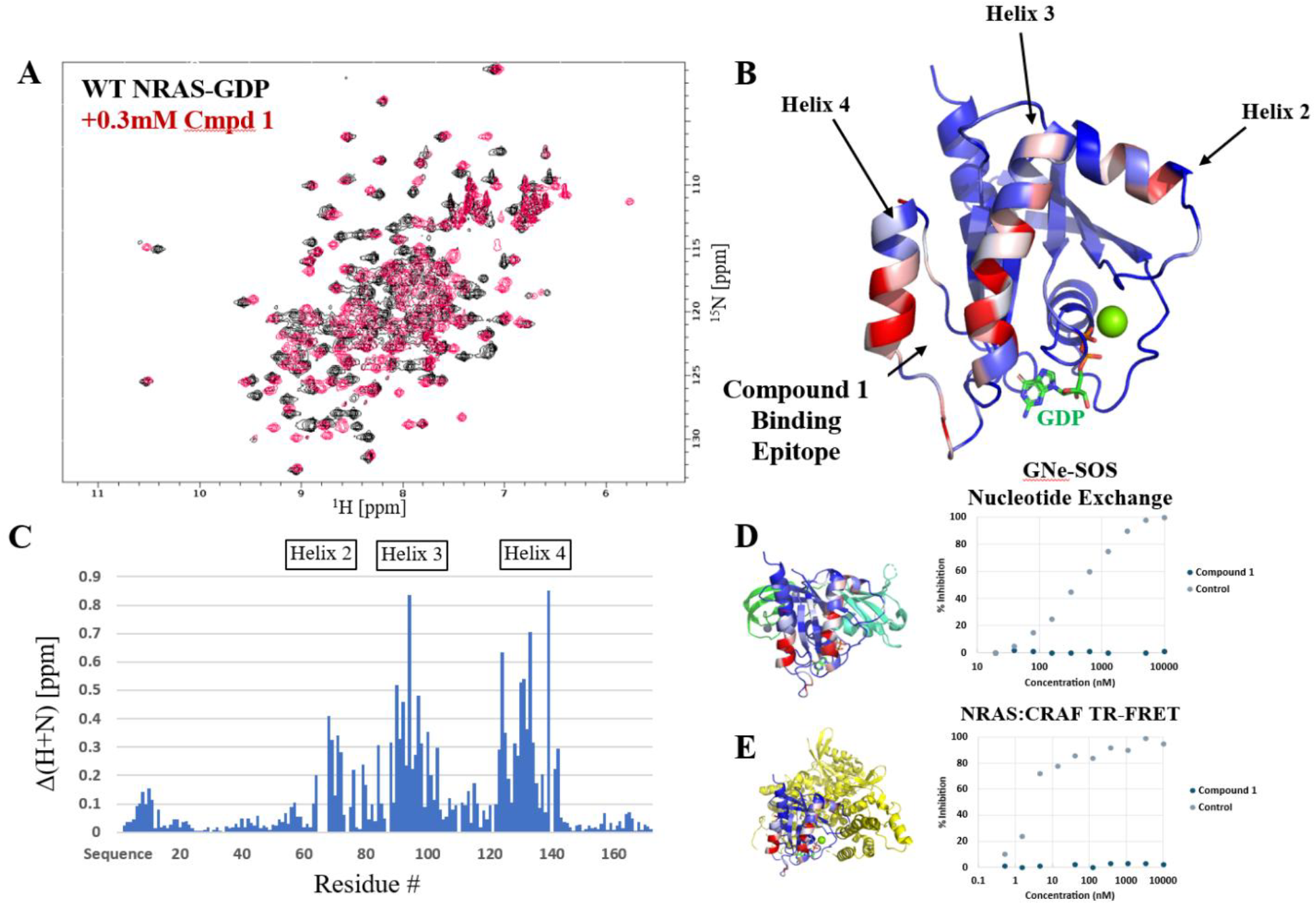
Binding Site and Function of Compound 1. A) 2D ^1^H/^15^N HSQC of apo WT NRAS (Black) and bound to 0.3 mM Compound 1 (Red). B) 3D Structure of NRAS with CSP of Compound 1 binding mapped in color (red most shifted, blue least shifted). C) CSP shift +/-Compound 1 binding plotted by WT NRAS residue number. D) GNE-SOS ATP transfer assay data for Compound 1 compared to Compound 40, an analog of RAS inhibitor LUNA-18 [20]. SOS protein is shown in Green & Teal. E) Fluorescence Polarization assay measuring the NRAS (Blue/Red) : CRAF (Yellow) protein-protein interaction for Compound 1 compared to control Compound 40 [20].

Combining the beneficial d-Pro mutation with other size-reduction changes led to Compound 25, a quadruple mutant with the sequence ClAcNle-PyD-Tic-Y-NMeF-dP-Bip-P-F-N-C. Compound 25 has only 8 Hydrogen bond donors, 5 fewer than Compound 1, while retaining an affinity of 1.1 µM for WT NRAS-GDP (Table 1). However, neither Compound 1 nor Compound 25 has detectable passive permeability as assessed by a double-sink parallel artificial membrane permeability assay (Figure S2). Given the molecular weight of the two compounds remains >1500 Da (Fig S3), the permeability result is unsurprising despite the reduction in polarity.

Affinity for RAS-OFF (loaded with GDP) and RAS-ON (loaded with GTP analog GMP PNP) NRAS changed together for 19 of the 25 compounds (Fig 3). Some peptides with changes to position 2, 6 and 8 diverge, suggesting part of the peptide:protein interface is influenced by the GTP-binding site. Compound 7 (Position 2, D to SerOMe), slightly improved affinity for RAS-ON NRAS while reducing affinity for RAS-OFF NRAS. Alternatively, installing A at position 2 (Compounds 9, 22) or NMeA at position 6 (Compound 13, 21) selectively breaks binding to one form or the other to achieve activation-state selectivity. Compound 22 is particularly interesting, as it is a 500 nM binder to RAS-ON NRAS with >10 µM affinity for RAS-OFF NRAS, suggesting it may be possible to specifically target RAS-ON NRAS at the helix 3-helix 4 interface.

**Figure 3:**
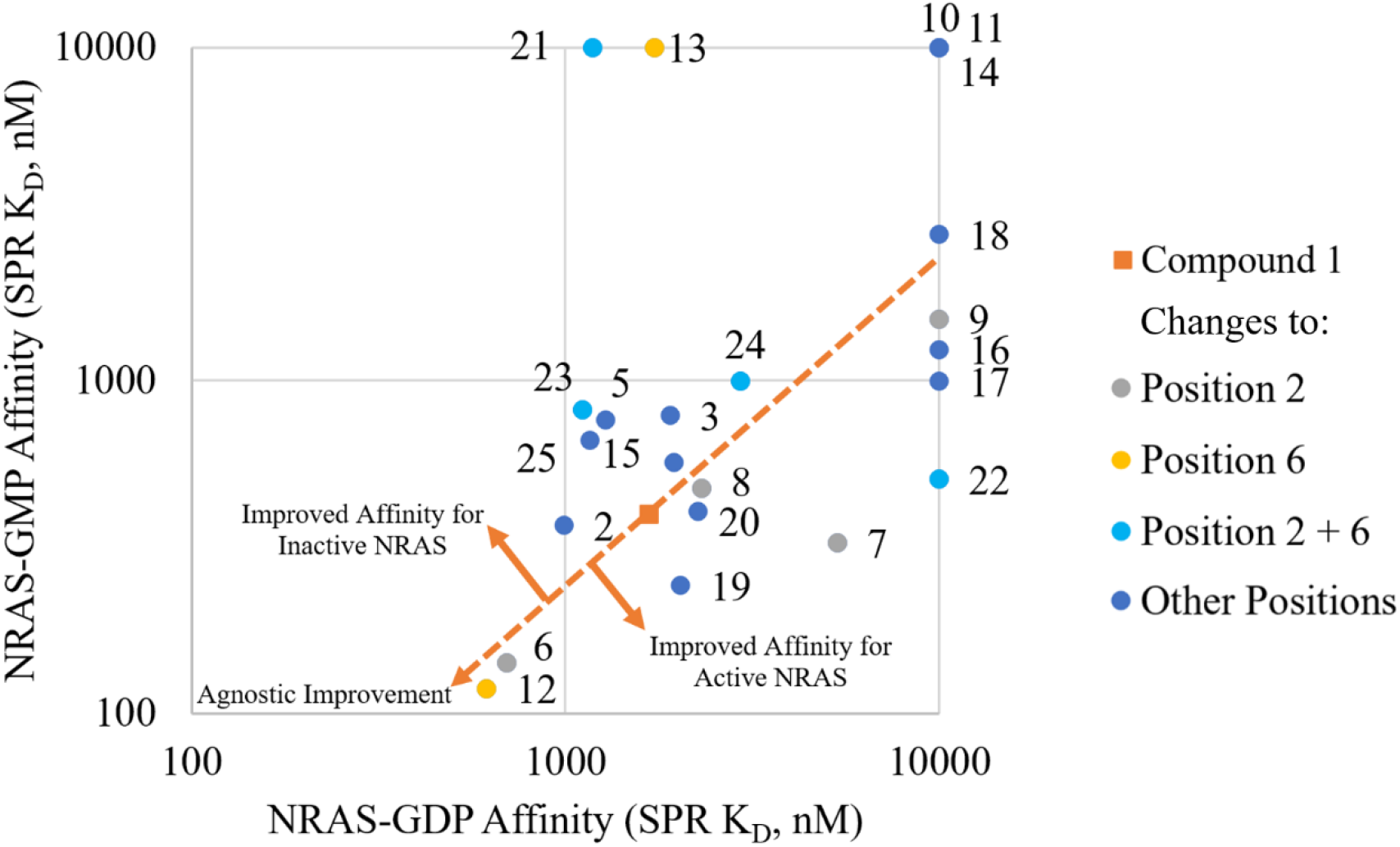
Selective Affinity for RAS N or RAS FF NRAS. Compounds 1-25 plotted by their SPR affinity to inactive, RAS-OFF NRAS (loaded with GDP) or activated, RAS-ON NRAS (loaded with GMP-PMP). The screening hit Compound 1 is in orange, with other compounds plotted in blue, light blue, or yellow depending on the position changed in variation from Compound 1. The dotted orange line represents equal potency for RAS-OFF and RAS-ON NRAS. To visualize non-binders, their markers are placed at the 10,000 nM line. The full SPR data is in Table S2.

All variants - regardless of affinity or state-selectivity - failed to inhibit NRAS in biochemical activity assay (Table S1). However, the Compound 1 site was recently implicated in NRAS homodimerization, disruption of which is a potential allosteric inhibition mechanism [21]. We therefore tested Compound 1 for cellular activity by monitoring phosphorylation levels of ERK1/2 in HMVII (NRAS – Q61K) cells using an AlphaScreen. Compound 1 had no pERK inhibition either with or without digitonin permeabilization (Fig S3). While the Compound 1 family lacks biochemical and cellular potency, the SAR series does reveal the helix 3/4 binding site can be used to simultaneously encode NRAS isoform selectivity and RAS-ON or RAS-OFF selectivity into a single peptide.

## Conclusions

Discovering molecules which discriminate between highly similar proteins is challenging, but mRNA display can provide macrocyclic peptides with exquisite selectivity when screening is performed under conditions that drive enrichment of specific binders. Here we use mRNA display to discover Compound 1, a peptide which engages a novel epitope on the allosteric lobe of the NRAS G-domain to achieve selectivity over KRAS and HRAS. While Compound 1 is small enough to have the potential for passive permeability, our efforts to optimize the sequence did not lead to a cell-permeable compound, emphasizing the ongoing challenge of encoding cell permeability into peptides of this size that have been selected for affinity. Further investment, such as structure-based drug design, could yield a permeable variant.

In addition to isoform selectivity, Compound 1 binds to both WT and cancer mutant NRAS (Q61R & Q61K) and displays a 10-fold affinity preference for the NRAS that has been activated by loading with GTP analog GMP-PNP. Single amino acid changes to Compound 1 can tune its affinity to prefer RAS-ON or RAS-OFF, confirming the allosteric coupling between the Compound 1 binding site and NRAS activation state. This connection deserves further study, as an understanding of the molecular mechanism of this family’s activation-state selectivity could reveal a new mechanism for targeting the oncogenic NRAS-ON state. Given the lack of biochemical inhibition for all Compound 1 analogs we designed – even those with SPR affinities of <200 nM – it is unlikely that the site can be used to achieve inhibition directly. Instead, the discovery of Compound 1 demonstrates how mRNA display can be used to find ligands at new selective sites on highly conserved proteins and could be used to study of NRAS-specific biology or as a starting point for building RAS-ON selective heterobifunctional NRAS degraders.

### Experimental

#### Protein Constructs

Human NRAS (residues 2-172) variants were synthesized and subcloned into the pET28a vector (Novagen; Millipore Sigma) with a N-terminal 6xHis tag, TEV cleavage site, followed by a SUMO tag and Avi tag. An additional NRAS (2-172) construct was designed to remove the Avi tag leaving native NRAS amino acids for BioNMR studies. For selectivity studies, KRAS (2-16) and HRAS (2-166) were used. Additional purification details are available in the supplemental information.

#### mRNA Display

Peptide Discovery Platform System (PDPS) was established at Merck & Co., Inc., Rahway, NJ USA under license from PeptiDream (Kawasaki, Kanagawa, Japan). The methods described herein are based on previously published work and adjusted to reflect the experiments performed in this study [1]. Full experimental details are available in the supplemental information.

#### Affinity Selection

Saturating amounts of biotinylated NRAS were immobilized on M-280 magnetic beads (Dynabeads M-280 streptavidin; Thermo-Fisher) by incubating the target protein with beads for 20 minutes at 4 °C. Panning was carried out by resuspending NRAS-bound M-280 beads to a final concentration of 200 nM protein with desalted peptide-mRNA-cDNA libraries for 1 h at 4 °C with rotation. In round 2 of the selection, a KRAS subtraction step a was incorporated before the panning step to remove potential KRAS-binding peptides. After five rounds of selection, post-Round 5 cDNA were sequenced using a MiSeq next-generation sequencer (Illumina). Additional details are available in the supplemental information.

#### Peptide Solid Phase Synthesis

Peptides were synthesized on a Syro II synthesizer using standard protocols for the generation of macrocyclic peptides arising from cyclization of Cysteine or Cysteamine onto a N-terminal amino acid capped as chloro acetamide. Full synthetic workflow is provided in the supplemental information. Representative quality control data for 5 synthetic peptides are provided in Figure S3.

#### Parallel Artificial Membrane Permeability Assay

PAMPA permeability was assessed using 6-well Stirwell sandwich plates (Pion Inc., catalog #11024350). Samples were finally analyzed and quantified by liquid chromatography–mass spectrometry (LC-MS). Experimental and analysis setup is described in supplemental information.

#### Surface Plasmon Resonance

The SPR binding assays were performed on a Biacore™ S200 or an 8K+ SPR system (Cytiva) using Biotin CAPture Kit with Series S Sensor Chip CAP (Cytiva) following manufacturer recommendations. Capture of biotin-tagged Ras protein and regeneration of protein surface at the end of each peptide test cycle was enabled by use of a proprietary Biotin CAPture Reagent that includes streptavidin bound to single-stranded DNA (ss-DNA) that can hybridize to the complimentary ss-DNA strand on the CAP sensor chip. Additional experimental detail is available in the supplemental information.

#### NMR spectroscopy

NMR experiments were carried out on a Bruker NEO 600 MHz spectrometer equipped with 5-mm TCI cryoprobe. The two NMR samples are 0.25mM ^13^C/^15^N labeled wild-type NRas(1-172)-GDP, with or without 0.3 mM Compound 1 peptide, in a 50 mM HEPES, pH 7.4, 50 mM NaCl, 2.0 mM MgCl2, 100 µM 3-(Trimethylsilyl) propionic-2,2,3,3-d4 acid (TSP), and 10% (v/v) D2O buffer. All 3D experiments were collected at 2 8K with nonuniform sampling (NUS) at a sampling rate of 25%. All NMR data were processed on a Linux station using NMRPipe and the spectra were analyzed using NMRFAM Sparky. The backbone resonance assignments were achieved by analyzing 2D ^1^H/^15^N HSQC, and triple resonance HNCA, HN(CO)CA, HNCO, and HN(CA)CO. In total, 7% of ^1^H and ^15^N backbone resonance shifts (163 of 168 non-proline residues) were assigned of both the apo and the peptide bound NRas. The CSPs induced by the peptide binding were calculated using the following formula: Δδ(Compound 1) = (((Δδ_1H_)^2^ + (0.14 × Δδ_15N_)^2^) / 2) ^1/2^.

#### SOS-catalyzed nucleotide exchange assay

Recombinant NRAS-WT, KRAS-WT and HRAS-WT (residues 2-16) were used in a SOS-mediated nucleotide exchange assay. Specifically, the SOS-catalyzed nucleotide exchange assay utilized a preformed TR-FRET complex containing a specific biotinylated RAS protein (NRAS-WT, KRAS-WT or HRAS-WT) with Bodipy-GDP, and Terbium-streptavidin. Compounds were preincubated with this complex for 60 minutes. Subsequently, recombinant human SOS protein and unlabeled GTP are added to initiate the exchange reaction. Antgonist peptides stabilize the Bodipy-GDP complex whereas the untreated protein rapidly exchanges Bodipy-GDP for unlabeled GTP resulting in reduced TR-FRET signal. Assay details are available in the supplemental information.

#### C-RAF-RBD TR-FRET displacement assay

The RAF-Ras binding domain (RBD) protein interaction assay utilizes recombinant biotinylated RAS-WT (NRAS or KRAS) protein loaded with GMPPNP and the GST-tagged Ras binding domain of C-RAF. The assay plate was incubated at ambient temperature with gentle shaking for 60 minutes to allow the reaction to come to binding equilibrium. Plate reader and analysis information is available in the supplemental information.

#### Cellular Phospho-ERK Assay in HMVII (NRAS – Q61K) cells AlphaScreen

Cellular NRAS inhibitory activity was evaluated by phosphorylation levels of ERK1/2 in HMVII (NRAS – Q61K) cells. Cell culture details are available in the supplemental information. Phosphorylated ERK and total ERK levels were detected by Alpha SureFire Ultra Multiplex pERK kit (Revvity #MPSU-PTERK) using 5 µL acceptor bead mix and 5 µL donor bead mix, both prepared following the manufacturer’s protocol. Ratio of pERK vs total ERK in each well was used as the final readout. Dose response curves and EC50 were analyzed using a 4-parameter logistic equation in GraphPad Prism software (GraphPad, San Diego, CA).

## Data Availability

The data supporting this article have been included as part of the Supplementary Information.

## Conflicts of nterest

The authors declare no conflicts of interest.

## Acknowledgements

The authors are grateful for contributions of Zachary Brown, Samuel Taylor, and Erwin Abucayon to the synthesis and characterization of the peptides in this manuscript, and to Lisa O’Callaghan and Bahanu Habulihaz for guidance designing the PAMPA studies.

## Author Contributions

K.K.H wrote the manuscript, designed and performed the mRNA Display screening, and with H.J. designed the SAR series

H.J. and S.L. oversaw the synthesis of the peptides

Y.Z performed the NMR experiments

A.S and J.R performed the SPR experiments

T.M. designed and oversaw the preparation of the protein constructs

X.C performed and M.S.M. oversaw the biochemical activity assays

G.V. performed the cellular activity assays

K.K.H, P.H.C., and H.Y, performed and H.Z. oversaw the PAMPA assays

R.E.L, E.B.K., D.G.M, A.W. and S.A.S supervised the research

All authors interpreted the results, and revised and approved the manuscript

## References

[1] Prior IA, Hood FE, Hartley JL. The Frequency of Ras Mutations in Cancer. Cancer Res. 2020;80(14):2969–2974.

[2] Jänne PA, Riely GJ, Gadgeel SM, et al. Adagrasib in Non-Small-Cell Lung Cancer Harboring a KRASG12C Mutation. N Engl J Med. 2022;387(2):120–131.

[3] Skoulidis F, Li BT, Dy GK, et al. Sotorasib for Lung Cancers with KRAS p.G12C Mutation. N Engl J Med. 2021;384(25):2371–2381.

[4] DepMap, Broad (2025). DepMap Public 25Q2. depmap.org

[5] Nakamura, K., Ichise, H., Nakao, K. et al. Partial functional overlap of the three ras genes in mouse embryonic development. Oncogene 27, 2961–2968 (2008).

[6] Wasko UN, Jiang J, Dalton TC, et al. Tumour-selective activity of RAS-GTP inhibition in pancreatic cancer. Nature. 2024;629(8013):927–936.

[7] Cregg J, Edwards AV, Chang S, et al. Discovery of Daraxonrasib (RMC-6236), a Potent and Orally Bioavailable RAS(ON) Multi-selective, Noncovalent Tri-complex Inhibitor for the Treatment of Patients with Multiple RAS-Addicted Cancers. J Med Chem. 2025;68(6):6064–6083.

[8] Ostrem JM, Peters U, Sos ML, Wells JA, Shokat KM. K-Ras(G12C) inhibitors allosterically control GTP affinity and effector interactions. Nature. 2013;503(7477):548–551.

[9] Wang X, Allen S, Blake JF, et al. Identification of MRTX1133, a Noncovalent, Potent, and Selective KRASG12D Inhibitor. J Med Chem. 2022;65(4):3123–3133.

[10] Zhang Z, Guiley KZ, Shokat KM. Chemical acylation of an acquired serine suppresses oncogenic signaling of K-Ras(G12S). Nat Chem Biol. 2022;18(11):1177–1183.

[11] Whaby M, Ketavarapu G, Koide A, et al. Inhibition and degradation of NRAS with a pan-NRAS monobody. Oncogene. 2024;43(48):3489–3497.

[12] Zhang JZ, Li X, Batingana AR, et al. De novo design of Ras isoform selective binders. Preprint. bioRxiv. 2025;2024.08.29.610300.

[13] Tanada M, Tamiya M, Matsuo A, et al. Development of Orally Bioavailable Peptides Targeting an Intracellular Protein: From a Hit to a Clinical KRAS Inhibitor. J Am Chem Soc. 2023;145(30):16610–16620.

[14] Ohta A, Tanada M, Shinohara S, et al. Validation of a New Methodology to Create Oral Drugs beyond the Rule of 5 for Intracellular Tough Targets. J Am Chem Soc. 2023;145(44):24035–24051.

[15] Passioura T, Suga H. A RaPID way to discover nonstandard macrocyclic peptide modulators of drug targets. Chem Commun (Camb). 2017;53(12):1931–1940.

[16] Ishizawa T, Kawakami T, Reid PC, Murakami H. TRAP display: a high-speed selection method for the generation of functional polypeptides. J Am Chem Soc. 2013;135(14):5433–5440.

[17] Campbell S, Philips MR. Post-translational modification of RAS proteins. Curr. Op. Struct. Bio. 2021 (71): 180–192.

[18] Kessler D, Bergner A, Böttcher J, Fischer G, Döbel S, Hinkel M, Müllauer B, Weiss-Puxbaum A, McConnell DB. Drugging all RAS isoforms with one pocket. Future Med Chem. 2020 Nov;12(21):1911–1923.

[19] Walker ME, Zhu W, Peterson JH, et al. Antibacterial macrocyclic peptides reveal a distinct mode of BamA inhibition. Nat Commun. 2025;16(1):3395.

[20] US20240158446A1: Cyclic compound having selective inhibitory action on kras over hras and nras.

[21] Gebregiworgis T, Chan JYL, Kuntz DA, Privé GG, Marshall CB, Ikura M. Crystal structure of NRAS Q61K with a ligand-induced pocket near switch II. European Journal of Cell Biology. 2024; 103 (2):151412

