## Supplemental Information for "Discovery of an NRAS isoform and activation-state selective macrocyclic peptide"

### Supplementary Information

Figure S1 – Codon Table used in mRNA Display

| 1st | 2nd | | | | 3rd |
|---|---|---|---|---|---|
| | **U** | **C** | **A** | **G** | |
| **U** | **NMePhe** | Ser | Tyr | **Trp** | **U** |
| | | | | | **C** |
| | | | | **Cys** | **A** |
| | | | | | **G** |
| **C** | Leu | Pro | **Ahp** | Arg | **U** |
| | | | | | **C** |
| | | | | | **A** |
| | | | | | **G** |
| **A** | **NMeGly** | **Bph** | Asn | Ser | **U** |
| | | | | | **C** |
| | | | | | **A** |
| | **ClAcPhe** | | | | **G** |
| **G** | Val | **Tic** | Asp | Gly | **U** |
| | | | | | **C** |
| | | | | | **A** |
| | | | | | **G** |

Table S1: Biochemical Assay Data

| Sample | RAS GNE assay | | | | cRaf assay | |
|---|---|---|---|---|---|---|
| | NRAS EC50 (nM) | KRAS EC50 (nM) | HRAS EC50 (nM) | | NRAS WT GMPPNP EC50 (nM) | KRAS WT GMP PNP EC50 (nM) |
| Compound 40 [20] | 2.2 | 2.5 | 3.2 | | 66.1 | 541.7 |
| Compound 1 | >9975 | >9975 | >9975 | | >9975 | >9975 |
| Compound 2 | >9975 | >9975 | >9975 | | >9975 | >9975 |
| Compound 3 | >9975 | >9975 | >9975 | | >9975 | >9975 |
| Compound 4 | >9975 | >9975 | >9975 | | >9975 | >9975 |
| Compound 5 | >9975 | >9975 | >9975 | | >9975 | >9975 |
| Compound 6 | >9975 | >9975 | >9975 | | >9975 | >9975 |
| Compound 7 | >9975 | >9975 | >9975 | | >9975 | >9975 |
| Compound 8 | >9975 | >9975 | >9975 | | >9975 | >9975 |
| Compound 9 | >9975 | >9975 | >9975 | | >9975 | >9975 |
| Compound 10 | >9975 | >9975 | >9975 | | >9975 | >9975 |
| Compound 11 | >9975 | >9975 | >9975 | | >9975 | >9975 |
| Compound 12 | >9975 | >9975 | >9975 | | >9975 | >9975 |
| Compound 13 | >9975 | >9975 | >9975 | | >9975 | >9975 |
| Compound 14 | >9975 | >9975 | >9975 | | >9975 | >9975 |
| Compound 15 | >9975 | >9975 | >9975 | | >9975 | >9975 |
| Compound 16 | >9975 | >9975 | >9975 | | >9975 | >9975 |
| Compound 17 | >9975 | >9975 | >9975 | | >9975 | >9975 |
| Compound 18 | >9975 | >9975 | >9975 | | >9975 | >9975 |
| Compound 19 | >9975 | >9975 | >9975 | | >9975 | >9975 |
| Compound 20 | >9975 | >9975 | >9975 | | >9975 | >9975 |
| Compound 21 | >9975 | >9975 | >9975 | | >9975 | >9975 |
| Compound 22 | >9975 | >9975 | >9975 | | >9975 | >9975 |
| Compound 23 | >9975 | >9975 | >9975 | | >9975 | >9975 |
| Compound 24 | >9975 | >9975 | >9975 | | >9975 | >9975 |
| Compound 25 | >9975 | >9975 | >9975 | | >9975 | >9975 |

Figure S2: PAMPA Results

| Compound | Category | Pe (N= 4 avg) | % Recovery (N=4 avg) |
|---|---|---|---|
| Compound 1 | Parent Hit | 0 | 61 |
| Compound 21 | Derivative | 0 | 32 |
| Compound 22 | Derivative | 0 | 30 |
| Compound 23 | Derivative | 0 | 28 |
| Compound 24 | Derivative | 0 | 26 |
| Compound 25 | Derivative | 0 | 48 |

Figure S3: Synthetic Peptide Characterization, and representative LCMS Spectra

| **Sample** | Chemical Formula | Calculated Mass (M+H) | Observed Mass (M+H) | Mass (M+H) Error, ppm | Retention Time (s) |
|---|---|---|---|---|---|
| **Compound 1** | C87H99N13O15S | 1598.7183 | 1598.7197 | -5.03 | 2.84 |
| **Compound 2** | C85H94N14O18S | 1631.667 | 1631.6588 | 4.44 | 2.82 |
| **Compound 3** | C82H96N14O18S | 1597.6826 | 1597.6897 | 4.44 | 2.82 |
| **Compound 4** | C82H96N14O18S | 1597.6826 | 1597.6897 | -5.34 | 2.66 |
| **Compound 5** | C79H90N14O18S | 1555.6357 | 1555.6274 | 0.31 | 2.99 |
| **Compound 6** | C85H100N14O18S | 1637.7139 | 1637.7144 | 2.43 | 2.96 |
| **Compound 7** | C89H101N15O17S | 1684.7299 | 1684.734 | -2.23 | 2.93 |
| **Compound 8** | C85H96N14O17S | 1617.6877 | 1617.6841 | -2.23 | 2.88 |
| **Compound 9** | C85H96N14O17S | 1617.6877 | 1617.6841 | -4.72 | 2.9 |
| **Compound 10** | C84H94N14O16S | 1587.6771 | 1587.6696 | 6.12 | 2.67 |
| **Compound 11** | C80H92N14O18S | 1569.6513 | 1569.6609 | 4.71 | 3.18 |
| **Compound 12** | C85H93FN14O17S | 1633.6626 | 1633.6703 | -6.15 | 2.87 |
| **Compound 13** | C87H96N14O18S | 1657.6826 | 1657.6724 | -4.01 | 2.95 |
| **Compound 14** | C86H96N14O18S | 1645.6826 | 1645.676 | 2.96 | 2.68 |
| **Compound 15** | C79H89ClN14O18S | 1589.5967 | 1589.6014 | -1.48 | 2.81 |
| **Compound 16** | C84H92N14O18S | 1617.6513 | 1617.6489 | -4.01 | 2.9 |
| **Compound 17** | C86H96N14O18S | 1645.6826 | 1645.676 | -6.35 | 2.86 |
| **Compound 18** | C83H92N14O18S | 1605.6513 | 1605.6411 | 1.42 | 2.84 |
| **Compound 19** | C84H94N14O18S | 1619.667 | 1619.6693 | 6.56 | 3.05 |
| **Compound 20** | C85H94N14O17S | 1615.672 | 1615.6826 | 4.71 | 3.07 |
| **Compound 21** | C85H93FN14O17S | 1633.6626 | 1633.6703 | 0.88 | |
| **Compound 22** | C87H99N13O15S | 1598.7183 | 1598.7197 | 1.39 | |
| **Compound 23** | C83H92N12O14S | 1513.6655 | 1513.6676 | 0.56 | |
| **Compound 24** | C88H99N13O15S | 1610.7183 | 1610.7192 | 0.44 | |
| **Compound 25** | C88H99N13O14S | 1594.7233 | 1594.724 | 5.00 | |

Compound 21

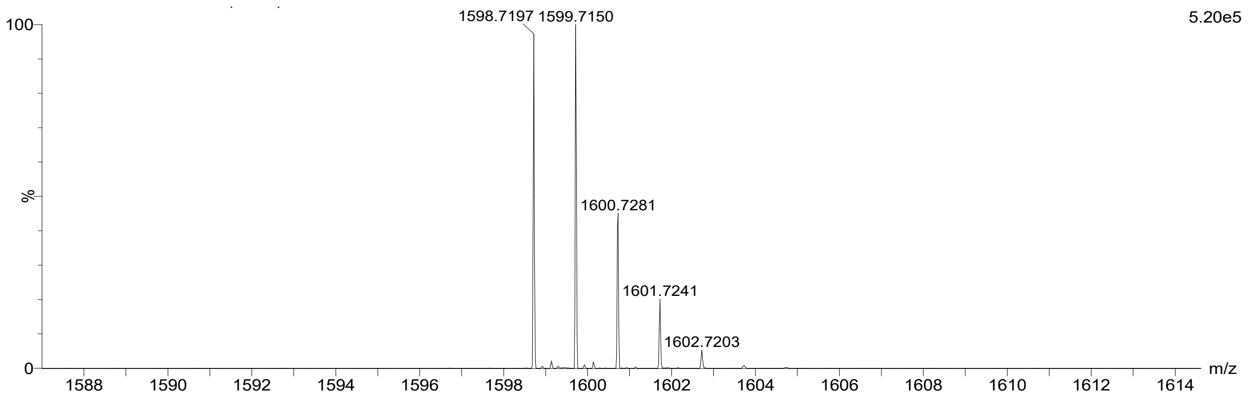

Compound 22

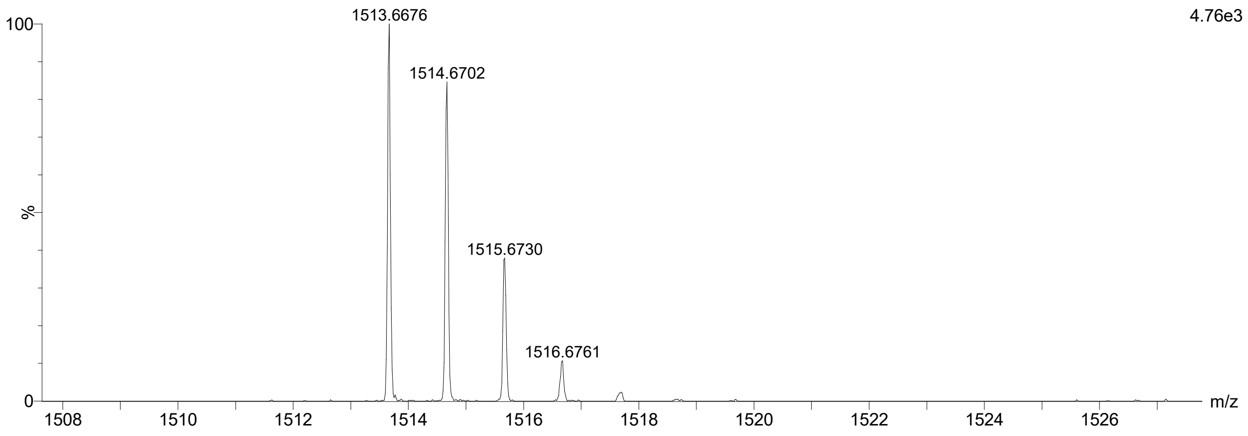


Compound 23

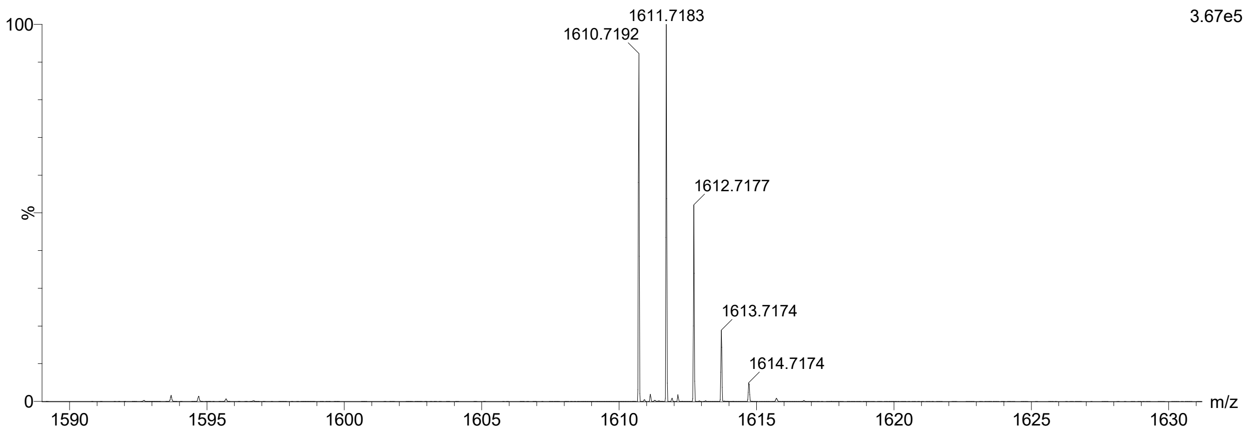


Compound 24

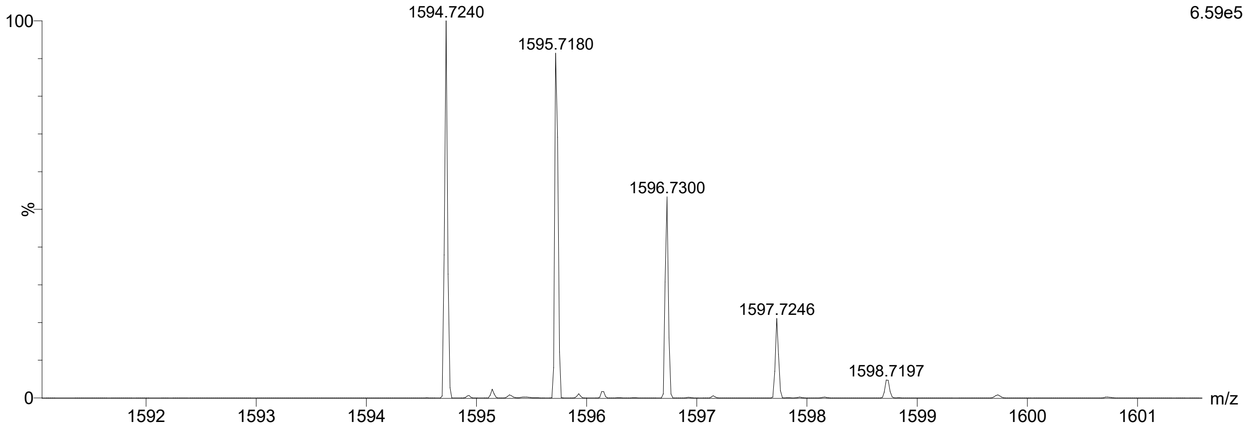

Table S2: Peptide affinities for all RAS constructs

| Compound ID | Sequence | WT KRAS-GDP | WT HRAS-GDP | WT NRAS-GDP | WT NRAS-GMP PNP | NRas Q61R-GTP KD (nM) | NRas Q61K-GTP KD (nM) |
|---|---|---|---|---|---|---|---|
| Compound 1 | **F-D-Tic-Y-NMeF-Sar-Bip-P-Y-N-C-G** | >10000 | >10000 | 1670 | 397 | 177 | 255 |
| Compound 2 | **Nle**-D-Tic-Y-NMeF-Sar-Bip-P-Y-N-C-G | >10000 | >10000 | 993 | 367 | 153 | 226 |
| Compound 3 | **L**-D-Tic-Y-NMeF-Sar-Bip-P-Y-N-C-G | >10000 | >10000 | 1910 | 786 | 380 | 495 |
| Compound 4 | **A**-D-Tic-Y-NMeF-Sar-Bip-P-Y-N-C-G | >10000 | >10000 | >10000 | 1570 | 408 | 662 |
| Compound 5 | **Cha**-D-Tic-Y-NMeF-Sar-Bip-P-Y-N-C-G | >10000 | >10000 | 1280 | 761 | 560 | 833 |
| Compound 6 | F-**PyD**-Tic-Y-NMeF-Sar-Bip-P-Y-N-C-G | >10000 | >10000 | 698 | 142 | 107 | 155 |
| Compound 7 | F-**SerOMe**-Tic-Y-NMeF-Sar-Bip-P-Y-N-C-G | >10000 | >10000 | 5340 | 325 | 376 | 477 |
| Compound 8 | F-**T**-Tic-Y-NMeF-Sar-Bip-P-Y-N-C-G | >10000 | >10000 | 2310 | 474 | 729 | 917 |
| Compound 9 | F-**A**-Tic-Y-NMeF-Sar-Bip-P-Y-N-C-G | >10000 | >10000 | >10000 | 1530 | 857 | 1260 |
| Compound 10 | F-D-**P**-Y-NMeF-Sar-Bip-P-Y-N-C-G | >10000 | >10000 | >10000 | >10000 | 1850 | 2560 |
| Compound 11 | F-D-Tic-**Phe4F**-NMeF-Sar-Bip-P-Y-N-C-G | >10000 | >10000 | >10000 | >10000 | 4810 | 6690 |
| Compound 12 | F-D-Tic-Y-NMeF-**dP**-Bip-P-Y-N-C-G | >10000 | >10000 | 615 | 119 | 32 | 47 |
| Compound 13 | F-D-Tic-Y-NMeF-**NMeA**-Bip-P-Y-N-C-G | >10000 | >10000 | 1730 | >10000 | 287 | 430 |
| Compound 14 | F-D-Tic-Y-NMeF-Sar-**Phe4Cl**-P-Y-N-C-G | >10000 | >10000 | >10000 | >10000 | >5000 | >5000 |
| Compound 15 | F-D-Tic-Y-NMeF-Sar-Bip-**Aze**-Y-N-C-G | >10000 | >10000 | 1950 | 567 | 381 | 539 |
| Compound 16 | F-D-Tic-Y-NMeF-Sar-Bip-**Pip**-Y-N-C-G | >10000 | >10000 | >10000 | 1240 | 496 | 754 |
| Compound 17 | F-D-Tic-Y-NMeF-Sar-Bip-**Sar**-Y-N-C-G | >10000 | >10000 | >10000 | 994 | 1110 | 1430 |
| Compound 18 | F-D-Tic-Y-NMeF-Sar-Bip-**NMeA**-Y-N-C-G | >10000 | >10000 | >10000 | 2740 | 2320 | 3480 |
| Compound 19 | F-D-Tic-Y-NMeF-Sar-Bip-P-**F**-N-C-G | >10000 | >10000 | 2030 | 243 | 167 | 225 |
| Compound 20 | F-D-Tic-Y-NMeF-Sar-Bip-P-**Phe4F**-N-C-G | >10000 | >10000 | 2260 | 406 | 480 | 625 |
| Compound 21 | F-**PyD**-Tic-Y-NMeF-**NMeA**--Bip-P-Y-N-C | >10000 | | 1180 | >10000 | | |
| Compound 22 | F-**A**-Tic-Y-NMeF-**dP**--Bip-P-Y-N-C | >10000 | | >10000 | 505 | | |
| Compound 23 | F-**PyD**-Tic-Y-NMeF-**dP**--Bip-P-Y-N-C | >10000 | | 1110 | 816 | | |
| Compound 24 | F-**PyD**-Tic-Y-NMeF-**dP**--Bip-P-**F**-N-C | >10000 | | 2930 | 995 | | |
| Compound 25 | **Nle**-**PyD**-Tic-Y-NMeF-**dP**--Bip-P-**F**-N-C | >10000 | | 1160 | 663 | | |

Figure S3: Compound 1 activity in pERK AlphaScreen in HMVII with NRAS Q61K

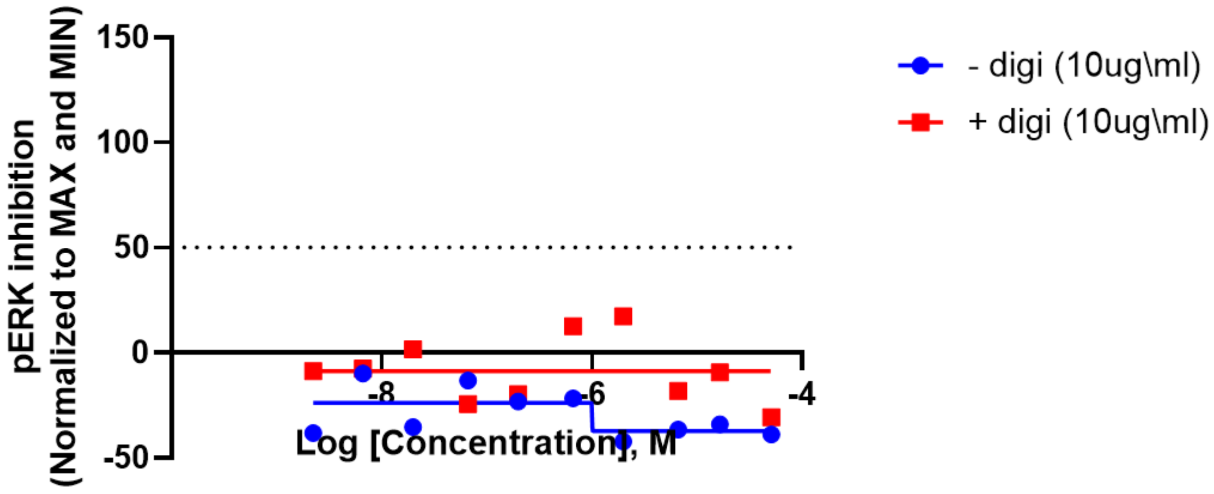

### Supplementary Experimental Details

#### *Protein Constructs*

In total, 4 NRAS constructs were used in this study: 6His-TEV-SUMO-Avi-NRas (2-172), 6His-TEV-SUMO-NRAS (2-172) 13C/15N, 6His-TEV-SUMO-Avi-NRas (2-172) Q61R and 6His-TEV-SUMO-Avi-NRas (2-172) Q61K. Sequences were obtained from UNIPROT: P01111 · RASN_HUMAN.

Plasmids were transformed into E.coli BL21(DE3) cells (NEB) followed by single colony selection on Luria-Bertani (LB) agar plates (Teknova) containing 50 ug/ml kanamycin. LB media (Teknova) containing 50 ug/ml kanamycin were inoculated with a single colony and grown overnight at 30°C. Overnight cultures were diluted and grown at 37°C until OD600 reached 0.8. Induction was initiated by adding 0.8 mM IPTG for 4 hrs until the cell density at OD600 reached 3.0. All Avi tag construct expressions were performed at 37°C in LB media. Expression of double labeled (U-15N, U-13C) NRAS (2-172) for BioNMR studies was carried out in M9 minimal media using 15N NH4Cl (1 g/l) and 13C glucose (4 g/l) as the nitrogen and carbon sources, respectively. BioExpress (U-15N, U-13C) at 0.1x concentration was added to media to boost expression levels. Isotopes were purchased from Cambridge Isotope Laboratories.

Induced cell pellets were resuspended and lysed in Buffer A; 40 mM HEPES pH 8.0, 300 mM NaCl, 20 mM imidazole, 2 mM beta-mercaptoethanol using Dounce homogenization followed by sonication (VibraCell VCX 750; Sonics). Cell debris was removed by centrifugation for 1 hr at 40,000 x g. Clarified lysates were applied to a His-Trap Crude Fast Flow column (Cytiva) and eluted with Buffer A including 500 mM imidazole. Peak fractions were pooled and desalted into Buffer A for SUMO protease cleavage reaction (1 mg SUMO protease/40 mg NRAS) carried out overnight at 4°C. NRAS was further purified using His-Trap Fast Flow to remove 6xHis-SUMO tag, SUMO protease and other contaminates in previously described buffers. Avi tag constructs were biotinylated using BirA ligase with reaction progression monitored using LCMS. Nucleotide exchange was necessary to remove GDP from wild-type NRAS and replaced with it non-hydrolyzable GTP analogue, GMP-PNP (Jena Bioscience). NRAS Q61 mutant constructs were preloaded with GTP making nucleotide exchange unnecessary. A Superdex-75 26/600 size-exclusion column (Cytiva) was used for the final purification step in 25 mM HEPES pH 7.5, 150 mM NaCl, 2 mM TCEP, 2 mM MgCl2, 5% glycerol. Monodispersed peak fractions were pooled and concentrated to ~2 mg/ml for biophysical characterization, BioNMR and screening activities.

#### *mRNA Display*

Partially reconstituted *E. coli*-based in vitro coupled transcription-translation systems were constructed by combining 50 mM HEPES-KOH (pH=7.6), 12 mM magnesium acetate, 100 mM potassium acetate, 2 mM spermidine, 20 mM creatine phosphate, 2 mM DTT, 2 mM ATP, 2 mM

GTP, 2 mM CTP, 2 mM UTP, 0.1 mM 10-formyl-5,6,7,8-tetrahydrofolic acid, 0.5 mM of each proteinogenic amino acid Leu, Val, Pro, Tyr, Asn, Asp, Arg, Ser, and Gly. The in vitro transcription-translation systems also contained 1.5 mg/mL total *E. coli* tRNA along with 0.73 µM AlaRS, 0.03 µM ArgRS, 0.38 µM AsnRS, 0.13 µM AspRS, 0.09 µM GlyRS, 0.02 µM HisRS, 0.4 µM IleRS, 0.04 µM LeuRS, 0.16 µM ProRS, 0.04 µM SerRS, 0.02 µM TyrRS, 0.02 µM ValRS, 2.7 µM IF1, 0.4 µM IF2, 1.5 µM IF3, 0.26 µM EF-G, 10 µM EF-Tu, 10 µM EF-Ts, 0.25 µM RF2, 0.17 µM RF3, 0.5 µM RRF, 0.1 µM T7 RNA polymerase, 4 µg/mL creatine kinase, 3 µg/mL myokinase, 0.1 µM pyrophosphatase, 0.1 µM nucleotide-diphosphatase kinase, and 1.2 µM ribosomes. The in vitro transcription-translation reaction also contained aminoacyl-tRNAs *N*-chloroacetyl-L-phenylalanine (ClAc-F)-$tRNA^{fMet}_{CAU}$, Cys-$tRNA^{Asn}$, MePhe-$tRNA^{Asn}$, MeGly-$tRNA^{Asn}$, Ahp-$tRNA^{Asn}$, and Bph-$tRNA^{Asn}$ and 2 µM library mRNA containing the open reading frame [5′-AUG-$(NNU)_{9\text{-}12}$-UGGGGAGGUGGUGGAAGUAGCUAG-3′] that was annealed to a puromycin linker. RF1 is omitted from the translation mixture to prevent multiple turnovers and to facilitate covalent attachment of the puromycin moiety to the C-terminal end of the GGGGSS peptide linker. After 30 min at 37° C, the 20 uL total volume transcription-translation reactions were halted by adding 20 µL 100 mM EDTA, pH 7.5 to dissociate the ribosomes from mRNA-peptide conjugates.

Reverse transcription of the peptide-tethered mRNA was carried out by adding a 40 µL solution containing 200 mM Tris, 300 mM KCl, 72 mM $MgCl_2$, 1.2 mM dNTPs, 12 µM Reverse Transcription Primer (5′-TAGCTACTTCCACCACCTCCCCA-3′), 800 units of M-MLV reverse transcriptase lacking RNase H activity (Promega, catalog# M368A) followed by incubation at 42 °C for 30 min. The reverse transcription reactions were quenched by adding 30 µL 100 mM EDTA, pH 7.5.

Macrocyclic peptide-mRNA-cDNA complexes were desalted using gel columns (Micro Bio-Spin P-6 Gel Columns, Bio Rad Cat#732–6221) that were pre-washed with selection buffer (25 mM HEPES pH 7.5, 150 mM NaCl, 2mM MgCl2, 2mM TCEP, 10 uM GDP), and the eluted library was adjusted to a final volume of 50 uL.

*Affinity Selection*

Additional method details:
After target incubation, the non-binding peptides were removed by washing the beads three times using selection buffer. The cDNA was eluted from the beads by incubating them at 95 °C for 5 min in polymerase-free PCR buffer (10 mM Tris-HCl pH 8.5, 50 mM KCl, 0.1% Triton X-100, 0.25 mM dNTPs, 2 mM $MgCl_2$, 0.25 µM Forward Primer (5′-CTAGTAATACGACTCACTATAGGGTTAACTTTAAGAAGGAGATATACATATG-3′), and 0.25 µM Reverse Primer (5′- CCCGCCTCCCGCCCCCCGTCCTAGCTACTTCCACCACCTCCCCA-3′) followed by collection of the supernatant. Ten units of Taq polymerase were added to the remaining supernatant and

the mixtures were amplified by PCR. Double-stranded DNA was transcribed into mRNA, which were standardized to 20 µM after quantitation using a Bioanalyzer 2100 (Agilent).

From round 2 onwards, a Kingfisher$^{TM}$ Purification System (ThermoFisher) was programmed to preclear peptide-cDNA-mRNA libraries by incubating them with KRAS loaded M-280 beads three times for 10 min before incubating them with NRAS-bound beads. In round 3 and all subsequent rounds, in vitro transcription-translation reactions were performed on a 5 µL scale, dsDNA PCR products from preceding rounds were added instead of mRNA, a puromycin linker was added directly to the reaction mixtures to anneal to in situ-transcribed mRNA.

*Peptide Solid Phase Synthesis*

The following steps were applied sequentially:

A) Peptide assembly: each peptide was synthesized on Rink amide resin on 25 umol scale. Fmoc-deprotection used 20% N-methylpiperidine in DMF (2 ml), mixing and stirring for 5 min followed by draining. The procedure was repeated once then each resin was washed with DMF (2 ml) four times. Each coupling involved Fmoc-amino acids (0.2 N in DMF, 8 equivalent), HATU (0.4 N in DMF, 8 equivalent), diisopropylethylamine (2 N, 16 equivalent), shaking for 30 min at 50 C, draining, washing with DMF (2 ml) twice. After the final amino acid coupling, each peptide was Fmoc-deprotected as described above, then coupled with chloro acetic anhydride (0.4N in DMF, 2 mL) and diisopropylethylamine in N-methyl pyrrolidone (2 N, 0.8 mL), shaking for 5 min, draining then the procedure was repeated once. Each resin was then washed with DMF (2 ml) twice, then DCM (2 mL) trice. B) Peptide cleavage: cleaving cocktail TFA:triisopropylsilane 96:4 (2 ml) was added to each resin followed by shaking for 1 h. Each mixture was drained into 50 mL centrifuge tubes filled with chilled diethyl ether:hexanes 4:1 (30 ml). Each resin was further washed with 0.5 ml of cleaving cocktail then centrifuged at 0 °C for 15 min. Each solution was decanted, each tube was filled with chilled diethyl ether (15 mL), shaken, then centrifuged at 0 °C for 15 min. Each solution was decanted, each tube was filled with chilled diethyl ether (15 mL), shaken, then centrifuged at 0 °C for 15 min. . The solutions were then decanted and the solids air dried for 2 h. C) Peptide cyclization: each solid was dissolved in DMSO (1.75 ml) at room temperature and treated with DIPEA (85 uL) then stirred for 2 h or longer depending on outcome as seen in LCMS. Each solution was then quenched with TFA (50 uL), filtered and directly purified on C8 column using a gradient of water:0.05% TFA in water to acetonitrile:0.05% TFA in water followed by evaporation.

*PAMPA and LC/MS analysis*

Up to four compounds were combined by mixing 5 µL of each 2 mM DMSO stock, generating a 500 µM mixture according to the sample sheet. From this mixture, 20 µL was diluted 100-fold with 1,980 µL of 1× PBS (pH 7.4) to yield an initial test concentration (Cd(0)) of 5 µM. To the diluted solution, 2 µL of a 1 mM verapamil stock (in PBS, pH 7.4) was added as an in-well control, resulting in a final verapamil concentration of 1 µM. The samples were mixed thoroughly and centrifuged at 13,000 rcf for 5 minutes to remove any precipitate.

Next, 200 µL of the clarified donor solution was transferred to the donor plate (bottom plate), and the remaining solution was retained as Cd(0) for recovery analysis. The acceptor plate (top plate) was inverted, and 5 µL of Pion's GIT-lipid solution was applied to the underside of each membrane support (avoid contact between the pipette tip and the membrane). After a 10-minute incubation, the acceptor plate was flipped upright and carefully aligned onto the donor plate, ensuring that no air bubbles were trapped. Then, 200 µL of Pion's acceptor sink buffer was added to the acceptor wells. The assembled sandwich plate was covered, placed in a sealed chamber with a moist paper towel to prevent evaporation, and the incubation start time was recorded.

Following the designated 12–16 hour incubation, the donor and acceptor plates were separated. From each donor well, acceptor well, and the Cd(0) solution, 25 µL aliquots were transferred into 96-well polypropylene storage plates for analysis. To match matrix conditions, 25 µL of acceptor sink buffer was added to donor and Cd(0) aliquots, and 25 µL of 1× PBS (pH 7.4) containing 1% DMSO was added to acceptor aliquots. All wells then received 200 µL of crash solution containing 250 ng/mL ITSD angiotensin II.

The incubated samples were analyzed on a Waters Xevo G2 XS quadrupole time of flight mass spectrometer (Waters, Milford, MA) coupled with Waters UPLC using a full scan acquisition method. Five µL of each sample were injected onto a 1.8 µm particle 100 × 2.1 mm id Acquity HSS T3 C18 column (Waters); the column was maintained at 55°C. The flow rate was 0.5 mL/min. Mobile phase A was water with 0.1% FA and mobile phase B was acetonitrile with 0.1% FA. The LC gradient started at 10% B for 0.5 min, ramped up to 98% B over 3 min, held at 98% B for 1 min before re-equilibration at 10% B for 0.5 min. The total run time was 5 min. The mass spectra were acquired in positive sensitivity mode using normal dynamic range, scanning in continuum mode from 300-2000 Da, at a scan rate of 0.2 sec ensuring that enough data points were acquired across each peak for robust quantitative analysis. The cone voltage was set to 35 V, with capillary voltage of 3.0 kV, source temperature of 125 °C, desolvation temperature of 450 °C, cone gas flow of 20 L/Hr, desolvation gas flow of 800 L/Hr. A single lock mass calibration at m/z 556.2771 in positive ion mode was used during acquisition to minimize possible drift in mass accuracy.

Targetlynx ® was used to preprocess the acquired mass spectra. The peak intensity and area of analyte peaks were exported as csv format and manually processed by Excel and Graphpad Prism software.

Angiotensin II human, Acetonitrile (Optima LC/MS grade), methanol (Optima LC/MS grade) and formic acid (FA, Optima LC/MS grade) were purchased from Fisher Chemical (Waltham, MA). Deionized (DI) water was purified with a Millipore system.

*Surface Plasmon Resonance*

Ras protein and test peptide working solutions were prepared in assay buffer consisting of 10 mM HEPES pH 7.4, 150 mM NaCl, 5 mM $MgCl_2$, 0.01% Tween-20, 2% DMSO, and supplemented with 5 or 10 µM nucleotide (GDP, GTP, or GMP-PNP) corresponding to the nucleotide bound to the Ras protein. Peptides were tested in a three-point concentration series (50, 200, and 1000 nM) or in a 6-point concentration series (6, 19, 75, 312, 1250, 5000 nM). Dilution series were prepared from peptide stocks (1, 2, or 10 mM in DMSO). Peptides were acoustically transferred to a 384-well plate using Echo 655T and backfilled with DMSO to 2 µl volume. Assay buffer (98 µL) was then added using a Bravo liquid handler to achieve final test concentrations at total volume of 100 µl. The final DMSO concentration was 2 %. SPR running buffer was prepared by adding DMSO to the assay buffer to a final concentration of 2%.

The SPR binding assay was run at 25℃. The sensor chip was loaded into the instrument and primed with running buffer. The sensor chip was first conditioned with three 60s pulses of regeneration solution (6M Guanidine-HCl, 0.25M NaOH). Biotin CAPture Reagent (diluted to 5 µg/mL in running buffer) was injected for 300s at 2 µL/min. Subsequently, biotin-Ras protein was immobilized to ~500-800 RU at a flow rate of 10 µL/min. Following a 60 s stabilization step with running buffer, test peptides were analyzed in single-cycle kinetic mode with an association time of 120s (20ul/min flow rate) and dissociation time of 600-900s. The sensor chip surface was then regenerated with a 120s-pulse at 10 µL/min of regeneration solution to start the next test cycle. Since DMSO was contained in the running buffer, a four-point solvent correction step was included at 0.375, 0.75, 1.5, and 3% DMSO to correct for any refractive index mismatches due to inherent inaccuracies in sample preparation and the hygroscopic nature of DMSO,
Data were collected with Biacore™ Insight Control software, and data were analyzed using the Biacore™ Insight Evaluation software and fit with the kinetic 1:1 binding model.

*SOS-catalyzed nucleotide exchange assay*

To assemble the preformed TR-FRET complexes, each biotinylated RAS protein is diluted to 2 µM in an EDTA Buffer (20 mM HEPES pH 7.5, 50 mM sodium chloride, 10 mM EDTA, and 0.01% Tween) and incubated at room temperature for one hour. This mixture is then further diluted to 90 nM in a buffer (20 mM HEPES pH 7.5, 150 mM sodium chloride, 10 mM magnesium chloride, and

0.005% Tween) containing 15 nM of Terbium-Streptavidin (Invitrogen, catalog# PV3577) and 900 nM of Bodipy-GDP (Invitrogen, catalog# G22360) and incubated at room temperature for six hours. It should be noted that this preformed TR-FRET complex for each of the RAS protein were made ahead of time, aliquoted and stored at -80 $^{o}$C until the day of the experiment.

Each test peptide (10 mM stock in DMSO) is dispensed into an assay plate (Corning, catalog#3820) using an Echo Acoustic Liquid Handler to make a final-10-point, ~3-fold dilution. Each well of the assay plate receives 5 µL RAS preformed TR-FRET complex diluted 45x in buffer containing 20 mM HEPES pH 7.5, 50 mM sodium chloride, 10 mM magnesium chloride, and 0.01% Tween) and is incubated at room temperature for 60 minutes (preincubation time). Each well then receives 5 µL of 2x recombinant human SOS protein and GTP (Sigma, G8877) in the same buffer and is incubated at room temperature for 60 minutes. The final reaction in each well of 10 µL consists of 3 mM GTP, 1 nM of specific Ras and 20 nM of SOS proteins.

The time-resolved fluorescence resonance energy transfer (TR-FRET) signal is measured on an Envision (PerkinElmer) plate reader: Excitation filter = 340 nm; emission1 = 495 nm; emission2 = 520 nm; dichroic mirror = D400/D505; delay time = 100 ms. The signal of each well is determined as the ratio of the emission at 520 nm to that at 495 nm. Percent effect of each well is determined after normalization to control wells containing DMSO (no effect) or a saturating concentration of inhibitor (max effect). The apparent effect as a function of compound concentration is fit to a four-parameter logistic equation.

*C-RAF-RBD TR-FRET displacement assay*

10 nL of each test peptide is dispensed into an assay plate (Corning, catalog#3820) using an Echo Acoustic Liquid Handler to make a 10-point, ~3-fold dilution. To the peptide assay plate, 5 µL of a 2x solution of biotinylated RAS-WT protein (GMPPNP-loaded; prepared in 20 mM HEPES 7.5, 150 mM NaCl, 10 mM MgCl2, 0.01% Tween 20) was added and incubated at ambient temperature for 30 minutes. Following the incubation, a 2x solution of GST-tagged C-RAF-RBD, anti-GST-d2 acceptor (Cisbio catalog #61GSTDLA) and Streptavidin-Tb cryptate donor (Cisbio catalog #610SATLA) prepared in the same assay was added to a final volume of 10 µL. The final concentrations in the assay are NRAS-GMPPNP:GST-C-RAF (2 nM:7.5nM) and KRAS-GMPPNP:GST-C-RAF (2 nM:50 nM), anti-GST-d2 (2 nM) and SA-Tb cryptate (25 nM).

The time-resolved fluorescence resonance energy transfer signal is measured on an Envision (PerkinElmer) plate reader with the following settings: dichroic mirror = LANCE/DELFIA DUAL/Bias; Emission1 = 615 nm; Emission2 = 665 nm; delay time = 60 ms. The TR-FRET signal from each well is determined as the ratio of the emission at 665 nm to that at 615 nm. Percent effect of each well is determined after normalization to control wells containing DMSO (no effect) or a saturating concentration of an antagonist control peptide (max effect). The percent effect as a function of compound concentration is fit to a four-parameter logistic equation.

*Cellular Phospho-ERK Assay in HMVII (NRAS – Q61K) cells AlphaScreen*

HMVII cells (Sigma) were cultured in T175 flask in growth medium (RPMI 1640 Medium, GlutaMAX™ Supplement) supplemented with 10% fetal bovine serum (Hyclone SH30071.03) and 1X Penicillin/Streptomycin (Gibco 15140-122). The cells were harvested in seeding medium (RPMI 1640 Medium, no phenol red (Gibco 11835-030) supplemented with 10% fetal bovine serum (Hyclone SH30071.03) and 1X Penicillin/Streptomycin (Gibco 15140-122) after 5 min of 0.25% Trypsin-EDTA (Gibco 25200-056) digestion and were seeded in 384-well tissue culture treated plate (Greiner 781091) at a density of 15,000 cells/20µL/well, and incubated at 37°C, 5% CO2 overnight. Prior to dosing of peptides, seeding medium was removed using the BlueCatBio Bluewasher system and replaced with 20 µL of assay medium without fetal bovine serum (RPMI 1640 Medium, no phenol red (Gibco 11835-030) and 1X Penicillin/Streptomycin (Gibco 15140-122) in the presence/absence of digitonin (10 µg\ml) 2 hours. For assays performed in the presence of 10% serum, seeding medium was used instead of assay medium. The compound dose-response titrations were prepared, and appropriate amounts of compounds were dispensed into the 384-well cell culture assay plate using the Echo 550 liquid handler. 25 µL assay or seeding medium was added to achieve a final assay volume of 45 µL. Assay plate was incubated at 37°C, 5% CO2 for the indicated length of time. After treatment, 25 µL medium was removed and transferred to an empty 384-well tissue culture treated plate (Greiner 781091) for the CytoTox-ONE™ Homogeneous Membrane Integrity (LDH) Assay. Remaining medium was removed from the plate using the BlueCatBio Bluewasher system, and cells were washed once with 25µL 1 × DPBS (Gibco 14190-144). Cells were lysed in 20 µL 1 × lysis buffer from Alpha SureFire® Ultra™ Multiplex pERK and total ERK assay kit (Revvity #MPSU-PTERK) containing EDTA-free Protease inhibitor cocktail (Roche 11836170001) at ambient temperature with constant shaking at 300 rpm for 10-15 min. The cell lysates were mixed for 10 cycles using the Agilent Bravo 384ST liquid handler system before 10 µL was transferred to OptiPlate-384 plate (PerkinElmer 6007680).

Plates were sealed using aluminum sealing tape (Costar 07-200-683) during incubation at ambient temperature with constant shaking at 300 rpm for 1 h (both acceptor and donor). Assay plates were read on a Envision Xcite Multilabel Reader (PerkinElmer #1040900) at ambient temperature, with emission at 535 nm (Total ERK) and emission at 615 nm (Phospho ERK).

CytoTox-ONE™ Homogeneous Membrane Integrity (LDH) Assay: Transient damage to cellular membranes were measured by LDH release assay. At 2hr post dose, 25 µL medium was removed from the cell culture assay plate and transferred to an empty 384-well tissue culture treated plate (Greiner #781091) using the Agilent Bravo 384ST liquid handler system. CytoTox-ONE reaction mix was prepared from the CytoTox-ONE™ Homogeneous Membrane Integrity Assay Kit (Promega#G7891) according to manufacturer's protocol. 25 µL of CytoTox-ONE reaction mix was added to the assay plate containing 25 µL medium, and plate was sealed using aluminum sealing tape (Costar #07-200-683). Assay plate was incubated at ambient temperature with constant shaking at 300 rpm for 45 min. Assay plates were read at ambient temperature on the Tecan M1000, with excitation at 560 nm and emission at 590 nm. Dose response curves and EC50 were analyzed using a 4-parameter logistic equation in GraphPad Prism software (GraphPad, San Diego, CA).
